# Taking a semi-local dynamic snapshot as a possibility for local homing in initially naïve bumblebees

**DOI:** 10.1101/178988

**Authors:** Anne C. Lobecke, Roland Kern, Martin Egelhaaf

## Abstract

For central place foragers, such as bumblebees, it is essential to return reliably to their nest. Instead of just flying away to forage, bumblebees, leaving their inconspicuous nest hole for the first time, need to gather and learn sufficient information about the surroundings to allow them to return to the nest at the end of their trip. Therefore, we assume an intrinsic learning program that manifests itself in the flight structure immediately after leaving the nest for the first time.

In this study, we recorded and analysed the first outbound flight of individually marked naïve bumblebees in an indoor environment. We found characteristic loop-like features in the flight pattern that appear to be necessary for the bees to acquire environmental information and might be relevant for finding the nest hole after a foraging trip.

Despite common features in their spatio-temporal organisation, the first departure flights from the nest are characterised by a high level of variability in their loop-like flight structure across animals. Changes in turn direction of body orientation, for example, are distributed evenly across the entire area used for the flights without any systematic relation to the nest location. By considering the common flight motifs as well as this variability, we came to the hypothesis, that a kind of dynamic snapshot is taken during the early phase of departure flights in the close vicinity of the nest location. The quality of this snapshot is hypothesised to be ‘ tested’ during the later phases of the departure flights concerning its usefulness for local homing.

## Introduction

The necessity of finding a route between the nest and a feeding site characterises a bumblebee’s everyday life as well as that of other hymenopterans. Bumblebees hatch inside their nest. When they leave it to forage for the first time, they are completely naïve and unfamiliar with its surroundings. The nest hole of most ground nesting insects, such as bumblebees, are inconspicuous and hard to find for humans, which makes it even more impressive, that bumblebees find the nest entrance after returning from a foraging trip. To accomplish this challenging task, the insect is required to gather sufficient information about the surroundings of the nest hole, suggesting an intrinsic learning program. This learning program should manifest itself in the flight structure of the departure flights immediately after leaving the nest for the first time. However, such a program cannot be expected to be entirely static and stereotyped, as it needs to be somehow adjusted to the particular environmental situation. This situation is unpredictable for the bee when leaving the nest hole for the first time and may differ much, for instance, when the nest entrance is oriented horizontally or vertically, or when the vegetation next to it is tightly cluttered or, alternatively, only loosely scattered.

Characteristic flight patterns, commonly interpreted as learning flights, have been observed in bees and wasps when they are unfamiliar with the surroundings of a relevant place. Then they perform peculiar flight sequences after leaving this place, which have been concluded to help gathering visual information about the environment near this place. Previous studies describe such learning flights as distinct and relatively stereotyped movement patterns with several common flight motifs. For social wasps, flight manoeuvres of increasing arcs are characteristic (Zeil, 1992; Zeil, 1993a; Collett and Lehrer, 1993; Stürzl et al., 2016). Thereby the insects continually gain height and turn in such a way towards a pivoting point that they keep the retinal image of the goal in the ventral part of the fronto-lateral visual field (Collett and Zeil 1996, Zeil et al. 2006, Zeil et al 2009). Similar flight patterns were also described for honeybees when leaving a profitable food source, most of these flights containing a high amount of translational movement and a backing away from the target structure while facing it for a large proportion of time (Dittmar et al. 2010, Lehrer and Collett 1994, Braun et al. 2010). This behaviour, often termed turn-back-and-look behaviour, has been described by Lehrer (1991; 1993) for honeybees as part of an efficient navigation system. Bumblebee departure flights from their nest hole show a loop-like structure which differs from the arcing pattern of social wasps and honeybees (Philippides et al. 2013). Rather than performing a turn-back-and-look behaviour bumblebees make small excursions away from the nest and then fly back towards the nest region and look at it (Hempel de Ibarra et al. 2009; Philippides et al., 2013; Collett et al., 2013). These movement patterns might be part of an efficient navigation system in bumblebees that allows the insects to gather, learn and later retrieve the information in the vicinity of their nest relevant for finding the way back to it.

Local homing of flying hymenopterans is assumed to rely mainly on visual cues, such as the spatial constellation of conspicuous objects close to the goal or the skyline of the panorama surrounding it (Collett and Collett, 2002; Collett et al., 2006; Towne et al., 2017). Another visual cue exploited is optic flow: Since stereopsis is not feasible for insects in the spatial range relevant for local homing, they largely rely on visual information from retinal image displacements generated by their structured movements (Gibson, 1950; Gibson, 1979; Srinivasan, 1993; Egelhaaf, 2009; Dittmar et al., 2010). Translational movement causes close target structures, such as the nest hole at departure and objects close to it, to shift further across the retina than objects further away (Stürzl and Zeil, 2007), which provides the insect with depth information (Lehrer and Collett, 1994). The location of the nest hole in relation to surrounding environmental features, such as vegetation might thus be gathered and memorised in this way (Dittmar et al., 2010).

Despite all these studies, the features of the environment and the flight manoeuvres that are essential to find a way back to a specific place are not yet entirely clear. Furthermore, it is still an open question, whether the insects learn during the entire first departure flight or only during specific parts of it, e.g. when passing the place primarily in translational movement or at the end of an arc. Here, we address these still unresolved problems by analysing the spatio-temporal characteristics of departure flights of naïve bumblebees (*Bombus terrestris*) after they leave their nest the first time. Considering that returning safely and fast to the nest is essential for bumblebees, our analysis will rest on the assumption that learning behaviour is the outcome of dynamic interactions between innate behavioural learning routines and visual information about the environment, which is actively shaped by just this behaviour as a consequence of the closed action-perception loop. The intrinsic learning program is expected to manifest itself, at least in a given environment, by a flight strategy with clearly invariant behavioural motifs. Therefore, we searched for invariants across animals in the spatio-temporal characteristics of the flight pattern that allow us to pinpoint the intrinsic behavioural program.

Several studies on local homing concentrated on the organisation of departure flights of bumblebees in semi-natural settings (Hempel de Ibarra, et al. 2009; Collett et al. 2013; Philippides et al. 2013; Riabinina et al. 2014). Since in such environments the rich environmental information can hardly be controlled by the experimenter, we did our analysis under laboratory conditions, where the environment is under control by the experimenter. For the first time, we recorded outbound and inbound flight sequences of marked bumblebees that have been initially naïve with respect to the visual environment of their nest entrance in a systematic way, allowing us to analyse the process and development of learning. The present paper is the first of a series of papers, which analyse the entire progression of learning and the consequences for the spatio-temporal organisation of successful return flights to the nest after foraging trips. In this first paper of the series, we focus on the very first outbound flights of bumblebees that are entirely naïve regarding the specific environment in which they forage and attempt to answer the following questions: In which way is the intrinsic behavioural program affected by the specific spatial layout of the surroundings of the nest entrance? How stereotyped is the innate learning strategy and how variable and inter-individually different may the behaviour be while still ensuring homing success?

## Material and Methods

### Animals and Experimental Setup

We obtained commercial bumblebee hives of *Bombus terrestris* (Linnaeus), containing only a few individuals, from Koppert (Berkel en Rodenrijs, The Netherlands). The bee hive was kept within a cubic Perspex box (each side measuring 30 cm) covered with black cloth in a room with a 12/12-hour light-dark cycle. A Perspex tunnel connected the nest box to another box of the same size, where the animals were free to fly and had access to an artificial feeder. In the first day after their arrival the bees had the possibility to learn how to use the artificial feeder filled with a commercial sucrose solution from Koppert, which was one of five feeders used later in the experiments. After one or two days the feeder was removed for most of the time and only returned to prevent the animals from starving during phases where no experiments were performed. The bumblebees had *ad libitum* access to pollen, directly put into the nest box. From the Perspex tunnel between the boxes another tunnel section led the bumblebees to a PVC tube (inner diameter 20 mm) connected to a hole in the floor of the test arena (Fig. 1A).

**Fig. 1.**
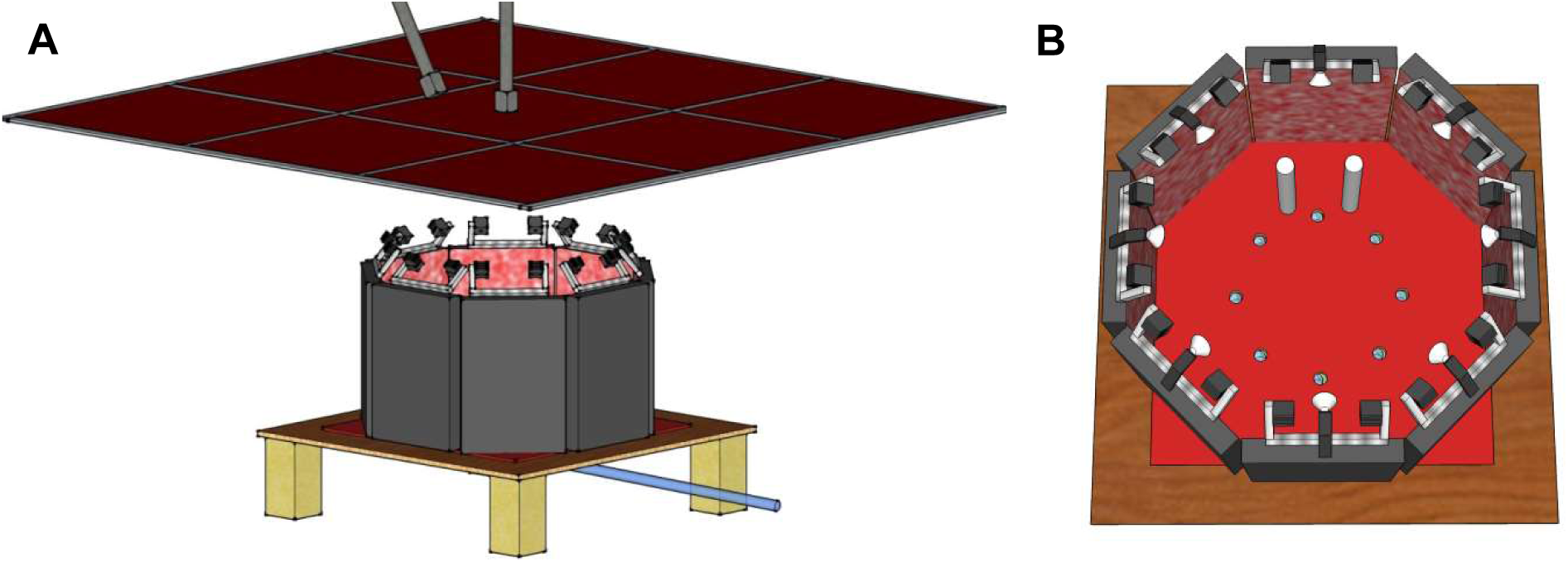
Experimental setup. (A) Flight arena seen from the side: red acrylic glass plate construction above the table with the flight arena. Grey structures above the glass plate construction are the high speed cameras. (B) Top view into the flight arena with 8 holes, two cylinders next to the hole that was connected to the nest; the other holes were closed some centimetres below the arena floor. The light setup consisted of 16 red LEDs (indicated by the grey boxes close to the left and right of each panel of the octagonal arena wall) and 8 white LEDs (indicated in white in the centre of each arena panel. The LEDs were mounted on the upper edge of the arena walls.

The behavioural analysis was performed in an octagonal test arena with an inner diameter of 95 cm, which was placed on a table (Fig.1A). Each wall segment was 60 cm high and 40 cm wide. The floor of the arena was covered with a red artificial grass carpet (Kunstgras Wereld, Antwerpen, Belgium) to add structure to the ground, but no distinct cues, ensuring a stable flight performance by the bumblebees. Eight holes (3 cm in diameter) were drilled into the arena floor. Each placed orthogonally to one of the wall segments at a distance of 22 cm (Fig.1B). Throughout the different experiments, only one of the eight holes was connected to the nest. The bumblebees could enter the arena via the PVC tube and started their flights from the connected nest hole. Two white cylinders were placed at a distance of 10 cm from that hole to indicate its connection to the nest. Apart from these cylinders, the nest hole could not be distinguished visually from the other holes. Regarding the holes in the floor, the arena was symmetrical and provided an ambiguous situation for the experiments. A 3 m * 3 m acrylic glass plate in red was mounted 40 cm above the arena (Fig.1A). Only light between 650 nm and 800 nm could pass through the acrylic glass. Therefore, the bumblebees, able to see light only up to 640 nm (Skorupski, 2010), were prevented from seeing the ceiling of the room and the cameras, which were placed above the glass plate (fig. 1b). To provide sufficient light for the camera recordings, eight white and 16 red LED lamps were positioned symmetrically with respect to the arena centre on top of its walls (Fig. 1B). The luminance inside the arena varied between 100 cd/m^2^ and 200 cd/m^2^

The bumblebees could leave the octagonal test arena into a large indoor flight room via the 40 cm gap between the arena walls and the acrylic glass plate. Beige curtains separated the flight area containing the test arena from the rest of the room. Ten fluorescent lamps (Biolux 965, Osram, Germany) illuminated the room (55 cd/m^2^ – 100 cd/m^2^). We used Biolux light with a spectrum between 400 nm and 700 nm to create as natural spectral lighting conditions as possible. In a corner of the flight room, bumblebees had access to feeders placed on a table. The bees were able to forage at those feeders and fly back to provide the hive with commercial sugar solution. In the experiments this ready-made solution was mixed with water at a ratio 3:1.

### Recording procedure

Bumblebees could be separated by removable doors in the tunnel system, so that only one bee at a time was allowed to enter the flight arena. Their outbound and inbound flights were recorded with two high speed cameras. These cameras (Falcon2 4M, Teledyne DALSA, Inc.) were placed above the acrylic glass plate (Fig.1A) and recorded the flights of the bumblebees at 148 fps, an exposure time of 1/1000 s and a resolution of 2048*2048 px. The optical axis of the top camera pointed straight downward. The optical axis of the second camera was 45° to the vertical. Using the software Marathon Pro by GS Vitec we recorded continuously for several hours on a hard disk array. Relevant sequences of outbound and inbound flights were stored as 8-bit jpeg images for the flight analyses. Sequences without relevant flights, i.e. where bumblebees just cross the recording area between upper walls and the acrylic glass plate construction, were deleted. A webcam (AXIS M10 Network Camera) was placed above the feeding table to monitor if bumblebees were foraging during the experiments.

### Training and Test procedure

The bumblebees entered the test arena through one of the nest holes in the arena floor. Only one of eight nest holes was connected to the nest during the experiments. We started the recordings immediately when we detected the bumblebee at the nest hole. During the training procedure, the two cylinders were placed next to the hole, which was connected to the nest and their positions were not changed during the first departing and return flights of each recorded bee. Bumblebees were able to forage at the feeding table during their flights in the flight room. After stopping a recording session at the end of one day, the end of the PVC tube leading to the arena was cleaned with 70% ethanol to remove potential odour cues placed by the bees. As a consequence of such a test arena, the space available for the bumblebees’ outbound and inbound flights was restricted. As an advantage of this restricted space we could record the entire flight and not only a few seconds or fragments of the flight. The flight structure obtained under these conditions does not differ in any obvious way from the departure flights obtained in other studies under different environmental conditions (Hempel de Ibarra, et al. 2009; Collett et al. 2013; Philippides et al. 2013; Riabinina et al. 2014).

### Data Analysis

The flight sequences from both cameras were analysed with the custom-built software ivTrace (Lindemann, 2005; <u>https://opensource.cit-ec.de/projects/ivtools</u>) where the position of the bee and the orientation of its body length axis were determined automatically. Additionally, ivTrace calculated the body orientation (yaw angle) from the top camera. In some cases, ivTrace had problems to track the elliptical form of the bumblebee’s body, and the yaw angle could not automatically be determined. This could happen when a bee crossed one of the nest holes or one of the edges between the wall segments. Then the software could only partially distinguish the bees from the dark background. In cases in which the automatic tracking procedure failed, the body position of the bee and the orientation of its length axis were determined manually. The Camera Calibration Toolbox of MATLAB (Jean-Yves Bouguet) was used for the camera calibration and the 3D stereo triangulation. A checkerboard pattern was used for the calibration. We determined the difference between recordings by the camera and the calculation.

The average position error for the top and the side camera were 0.11 px and 0.09 px, respectively. For some aspects of the analyses the bees’ the time-series of body orientation angles was filtered using a Gaussian filter with a window length of 1.35 ms. Besides the yaw angle of the bees’ body orientation several other parameters, e.g. height over ground and retinal position of the nest hole, were analysed and compared to characterise the flights’ spatio-temporal structure.

The analysis is based on 21 first departure flights of initially naïve bees with a total duration of 872 seconds.

## Results

This study is based on the assumption that the spatio-temporal organisation of outbound flights of bumblebees, after leaving the nest hole for the first time, is the outcome of dynamic interactions between innate behavioural learning routines and visual information about the environment. As a consequence of the closed action-perception loop, this information is actively shaped by the innate behaviour. The astonishing feat that a single departure flight in an unpredictable environment is sufficient for the initially naïve insects to find back to their home location, is worth to be investigated in a systematic way. That bumblebees and other hymenopterans gather relevant information about the environment on their departure flights from their nests is plausible as they perform peculiar flight sequences, and the departure flights decrease in duration and complexity with experience (Lehrer, 1991; Lehrer, 1993).

Here, we analyse for the first time systematically what is special about the structure of the first departure flight of naïve bumblebees, inter-individually and compared to other flying hymenopterans. Are there invariant motifs in the flight manoeuvres, which might be necessary for learning the location of the nest hole? To find this out we took a closer look on the flight structure of initially naïve bumblebees in an indoor test arena.

### Description of the overall flight structure

We observed a broadly similar flight pattern in bumblebees as described for social wasps and honeybees (cf. Introduction): The flights, starting from the nest hole, increased in height and distance to the starting point over time (Fig. 2). In contrast to the arcs of social wasps and the backing away from the nest hole of honeybees, the bumblebees performed loop-like excursions away from the nest and then flew back towards the nest region, a performance that is reflected in fluctuations of flight height and distance to the nest hole (Fig. 3). This flight characteristic is in accordance with what has been described for bumblebees under outdoor conditions (Hempel de Ibarra, et al. 2009; Collett et al. 2013; Philippides et al., 2013; Riabinina et al. 2014). In large parts of these loops, bumblebees faced towards the nest region (Fig. 4) as do wasps and honeybees for most of the time during the initial sections of their departure flights from the nest hole (Zeil, 1992; Stürzl et al., 2016; Collett and Lehrer, 1993).

**Fig. 2.**
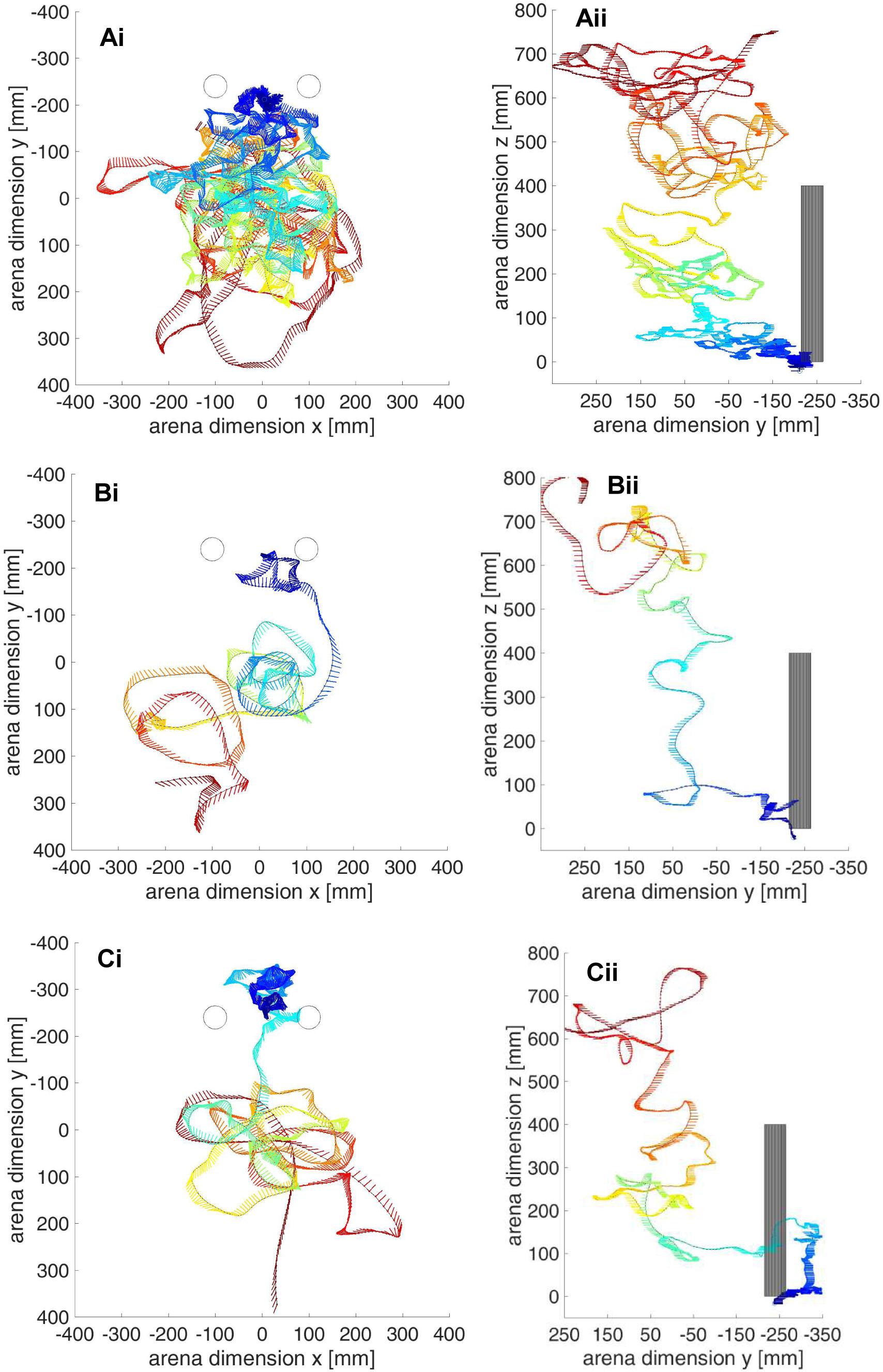
Flight trajectories of first flights of three different bumblebees seen from above and from one side. Black circles in the top view and grey rectangles in the side view: cylinders; blue circle in top view: hole connected to the nest; lines show the bee’s body orientation every 20.27 ms where the head’s position is on the trajectory. Flight trajectories colour-coded with time: Dark blue beginning of flight, dark red end of flight. Axes scales are given in mm. ‘ 0’ of the X- and Y-axis represents the centre of the flight arena

**Fig. 3.**
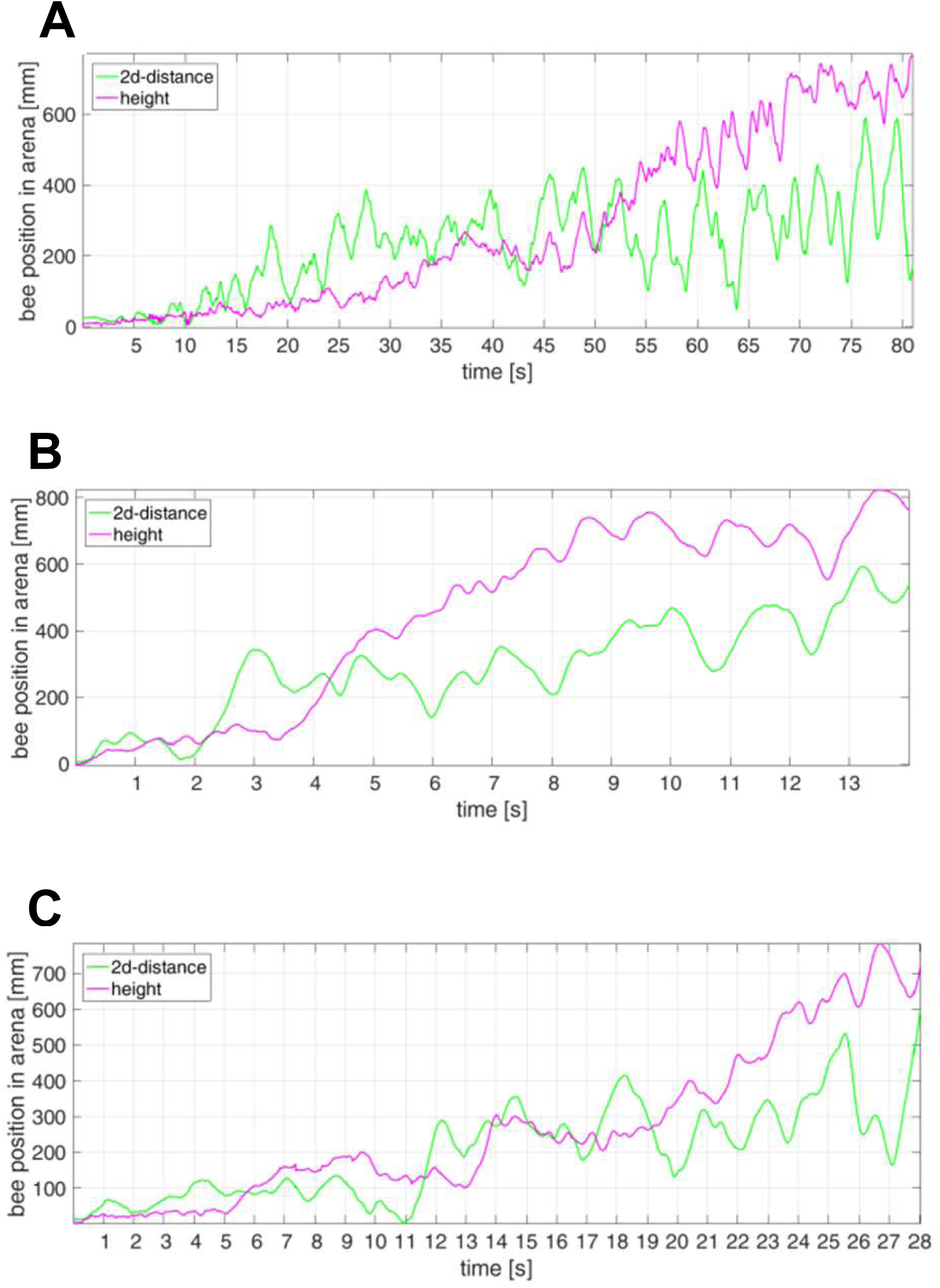
Time course of altitude and distance to the nest hole. Data is shown for the same three initial departure flights as shown in Fig. 2.

**Fig. 4.**
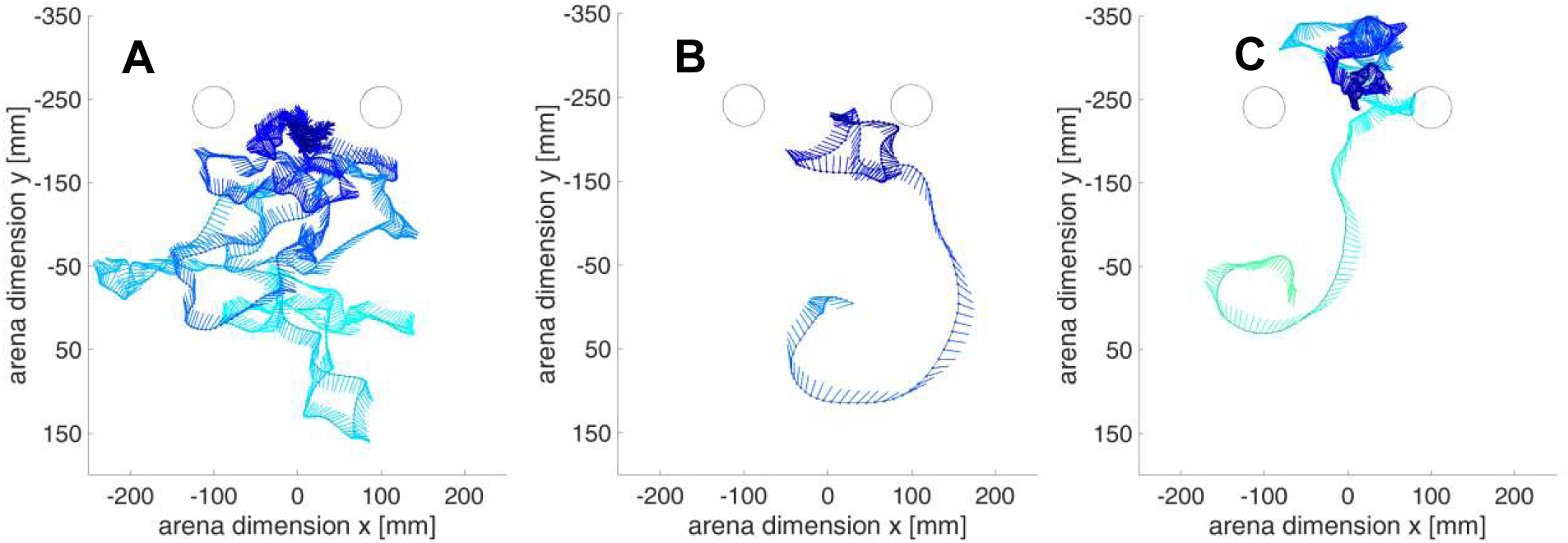
First phase of departure flights. Top view of Initial segments of the same three departure flights as shown in fig.2 for flight height above ground below 100 mm (seen from above). Lines show the bee’s body orientation every 20.27 ms where the head’s position is on the trajectory. Flight trajectories colour-coded with time as in fig. 2. The flight examples show many segments of translational movement.

After spending quite some time close to the nest hole, the bumblebees extended their departure flights towards the centre of the arena, where more space is available for their flights. The area between the nest hole and the closest arena wall was mostly avoided by the bumblebees. This suggests that they familiarise themselves with the immediate surroundings of the nest hole during this early part of the flight. Most of the time during this initial flight section, bumblebees flew close to the ground with an altitude roughly below 100 mm. After some time, they increased height and distance to the nest hole in loop-like flight patterns covering large parts of the horizontal extent of the flight arena including the nest hole region. When the bumblebees reached the height of the cylinders’ upper edge at 400 mm, they mostly circled around at this altitude, using the entire arena space.

These observations and previous studies suggest that learning of the nest hole location and its immediate environment occurs during the initial phase of the departure flights. Therefore, we decided to divide the flights into three different phases:

Phase 1 represents the part of the flights below 100 mm above ground level of the arena. This phase may include fluctuations in altitude where the bees’ altitude briefly exceeds 100 mm, but then returns to an altitude of less than 100 mm.

Phase 2 includes the flight sections between 100 mm and 400 mm altitude, excluding the brief flight sections where altitudes exceeded 100 mm (contained in phase 1) and including brief flight sections where the bees’ altitude briefly exceeds 400 mm, but then returns to an altitude of less than 400 mm. Phase 3 contains flight sections exceeding 400 mm altitude. Fluctuations which belong to phase 2 were excluded.

The thresholds do not represent altitudes with an obvious change in flight style and might, to some extent, be arbitrary. We ensured that the conclusions we will draw from our experiments are independent from the specific classification into the three flight phases.

### Leaving direction from the nest hole

When bumblebees leave their nest hole for the first time they do not know anything about its specific surroundings. This means that they cannot know in which direction to head for their search for potential feeding sites. Accordingly, the direction of the first departure from the nest hole should be arbitrary, unless the tube leading the bee to the nest hole were in some way asymmetric. Therefore, we analysed whether potential tube asymmetries affected the leaving direction of bees from the nest hole. This was done by subdividing the arena floor around the nest hole into eight 45°-segments and counting the bees entering each segment after leaving the nest hole. Only the first entered segment was counted, independent of the segment where the bumblebee started its flight. A Chi^2^-test showed no significant deviation from a uniform distribution at a significance level of p = 0.05 and, thus, no evidence that the tube properties influence the bumblebees’ direction of departure. A similar result was obtained for the direction of take-off around the nest location (Chi^2^-test, p = 0.05 significance level). These results suggest that the asymmetry in the flight pattern of the population of outbound flights (see next paragraph) is independent of the asymmetries in the tube system that leads the bees to the nest hole. Accordingly, the asymmetry in the overall flight pattern of all tested bees was most likely caused by the spatial layout of the test arena (i.e. location of cylinders and walls of the arena).

### Asymmetry of flight around nest hole

In our experimental setup bumblebees were confronted with an unpredictable environmental situation, including unequal distances to the eight wall segments of the arena and the two cylinders, which we positioned next to the nest hole. As long as the bees do not take into account any environmental information when shaping their flights, the overall distribution of flight paths across bumblebees should be symmetrical around the nest hole, because there is no need to prefer one direction, although individual flights might be asymmetric. Hence, as soon as asymmetries in the overall flight patterns across flights can be detected, spatial information about the surroundings of the nest hole is used by the bees to organise their flights. As figures 2 and 4 show, the bumblebees’ flights shifted towards the centre of the arena after an initial flight phase close to the nest hole. To find out when after flight onset spatial information is employed by the bees, we scrutinised the flight trajectories in two ways: To test whether the closest wall influenced the shape of the bumblebees’ flights, we first conceptually divided the arena by a horizontal line crossing the nest hole. This line served as a symmetry line for the flight pattern. The range closer to the wall was defined as range 1 and the one towards the centre of the arena as range 2 (fig. 5A). We expected the bees to spend more time of their flight in range 2, which is the direction of the centre of the arena, where more space is available. The time point when the bees started spending more time in range 2 rather than in range 1 is interpreted as the time point when the spatial layout of the arena plays a role in shaping the flights. On average, the bumblebees never tended to spend more time (over 50%) of their flights in range 1 rather than in range 2. After seven seconds of the flight they spend more than 75% of their flight on average in Range 2, the direction of the centre of the arena (fig. 5B).

**Fig. 5.**
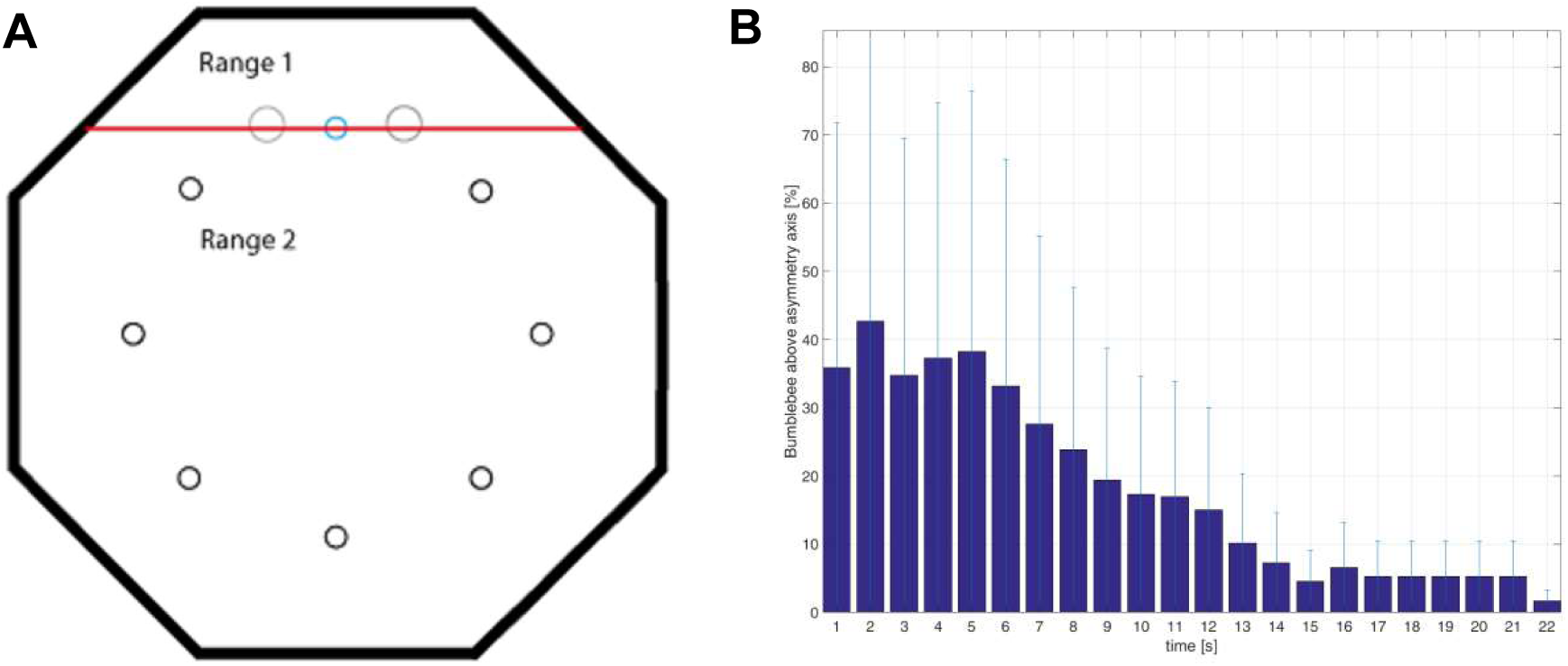
Asymmetry of flight around nest hole. (A) the initial departure flights. Arena divided into range 1 and range 2 (red line)Black circles: “Dummy” nest holes, Blue circle: Connected nest hole, Grey circles: Cylinders, (B) Percentage of time bumblebees spend in range 1 as defined in A as a function of time. For this analysis, time was binned in 1 s intervals. Dark blue bars: mean across bees, light blue: standard deviation, n = 21 flights

To directly test, whether this shift of the flight trajectories towards the centre of the arena is a consequence of the unequal distances to the eight wall segments, we did further experiments where we closed all eight peripheral nest holes and opened one nest hole in the centre of the arena, so that all wall segments were at the same distance to the nest hole and the flight structure should not have depended on the arena architecture. Now both ranges covered the same size of the arena: range 1 was above the horizontal line crossing the nest hole in the centre while range 2 was beneath it. Although individual flights observed under this condition (n = 8) were still asymmetrical and tended to cover one range of the arena, the outbound flights, on the whole, show no preference of one range over the other (data not shown, P = 0.0625). Another observation during these control experiments was, that, individual bees, after they started flying into a given range of the arena, stuck to it until they reached the height of the cylinders (400 nm), and then tended to use the whole arena for the last flight phase before leaving the arena. Still, across bees both ranges where chosen with the same likelihood.

We used the same flight data to test whether and after what time interval the two cylinders close to the nest hole shape the flight trajectories. Two conceptually perpendicular lines across the arena divided the space into four segments, of which two include a cylinder (fig. 6A). The analyses showed that the bumblebees avoided the segments containing the cylinders during most of their flight time (over 50%). They spend more than 75% of their flight time on average in range 2 after eight seconds (fig. 6B). These results thus reveal that, after leaving the nest for the first time, the innate learning routines of bumblebees are modified immediately or at the latest after a few seconds by spatial information about the specific surroundings, most likely extracted from the retinal image changes actively generated by the behavioural routines.

**Fig. 6.**
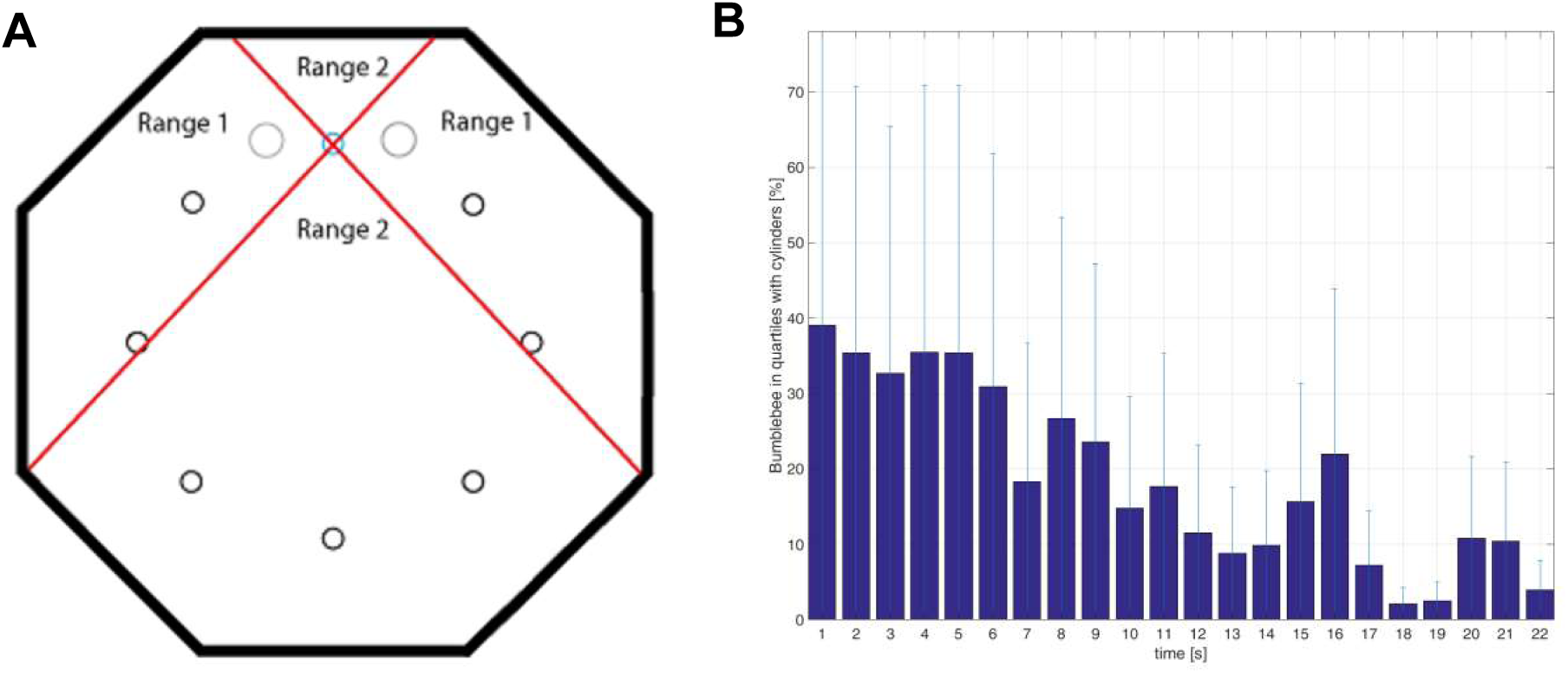
Asymmetry of flight around nest hole in respect to cylinders. (A) Arena divided into ranges 1 and range 2 (red lines). Black circles: “Dummy” nest holes. Blue circle: Connected nest hole. Grey circles: Cylinder. (B) Percentage of time bumblebees spend in range 1 as defined in A as a function of time. For this analysis, time was binned in 1 s intervals. Dark blue bars: mean across bees, light blue: standard deviation, N = 21 flights

### Turn-back-and-look behaviour – retinal position of the nest hole

Honeybees perform a so-called turn-back-and-look behaviour, where the bees turn around immediately after leaving the hive and face its entrance during the initial sections of the departure flight (Lehrer, 1991; Lehrer, 1993). Likewise, social wasps keep the retinal image of the target in the ventral part of the fronto-lateral visual field during the initial phase of departure flights (Collett and Lehrer, 1993; Collett and Zeil, 1996; Zeil et al., 2006; Zeil et al., 2009). Nevertheless, fixation of the nest hole has been reported to be rather inaccurate, since the image of the nest hole is kept within a rather extended retinal area after the insect gained distance from the nest (Zeil, 1993a). These studies suggest that it might be useful, if not essential, for hymenopterans to look with the frontal part of their visual field at the nest hole and its surroundings at least in the initial sections of the first outbound flight.

To assess whether this also holds for bumblebees, i.e. whether they keep the retinal image of the nest hole in a specific range of the visual field during significant parts of the initial phase of the outbound flights, a histogram of the retinal nest hole position was determined. Figure 7A shows that the nest hole is broadly kept in the frontal visual field between -60° and +60° across tested bees for most of the time. However, there seems to be no distinct region of the eye where the bumblebees fixated their nest hole. Rather, bees tended to roughly look towards the nest hole and its neighbouring regions for most of the time during the initial phase of outbound flights. This characteristic does not hold if bees gained height during the subsequent flight phases. In phase 2, a Chi^2^-test (significance level of p = 0.05) showed no significant deviation from a uniform distribution (fig. 7B). Furthermore, in phase 3, the retinal image of the nest hole was in the rear part of the eye for more time than it was in the frontal visual field (fig. 7C). This might be a consequence of the structure of flight trajectories: At higher altitudes, bumblebees used more space of the arena and tended to fly in increasing loops. The time intervals, where the bees face the nest hole region are, therefore, shorter than the time where the nest whole is seen roughly in the lateral regions and the rear part of the visual field.

**Fig. 7.**
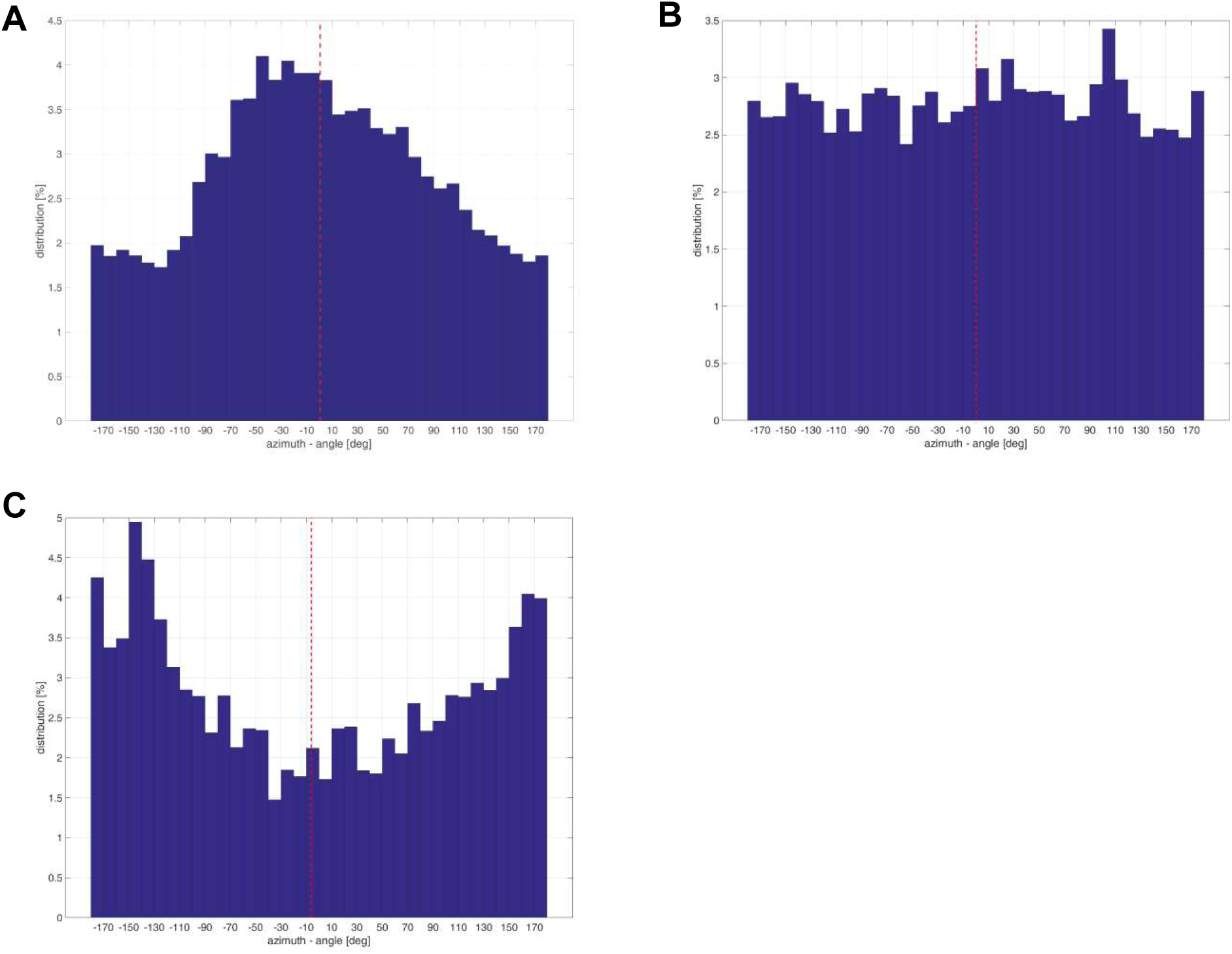
Histogram of the retinal nest hole position. (A) 1^st^ flight phase (below a height of 100 mm): the nest hole is broadly kept in the frontal visual field between -60° and +60° across tested bees for most of the time of a departure flight (B) 2^nd^ flight phase (height between 100 mm and 400 mm): no distinct region of the eye where the bumblebees fixated their nest hole (C) 3^rd^ flight phase (above a height of 400 mm): the retinal image of the nest hole was in the rear part of the eye for more time than it was in the frontal visual field: over 75% in -180° to -60° and 60° to 180°, but less than 25% in the region between -60° and +60° Red dashed line: mean of retinal position, N = 21 flights

Since the fixation of the nest hole in a broad frontal retinal area plays a significant role in the initial phase, we had a closer look at the first sections of the outbound flights. Zeil et al. (2009) observed that fixation periods in wasps occur during translations past the nest entrance, mostly during the arcs where the wasps tend to pivot around the nest entrance (Zeil et al., 2009; Boeddeker et al., 2010). To find whether this is a characteristic also of bumblebees’ first outbound flights we looked for locations in the flight arena, where the bumblebees kept the nest region in the frontal visual field between -25° and +25°. These locations are distributed throughout the whole area that is covered by the flight trajectories and do not correspond to distinct locations in the arena relative to the nest hole (fig. 8A-C). The duration of the flight sections while the bumblebees face the nest region varies for the individual bees as well as across bees and cover a broad range of time intervals (fig. 8D). Durations between 0 ms and 65 ms might be explained by a full rotation or loop flown by the bumblebee where the nest location crosses the insect’s retina between -25° and 25° necessarily. The other large portion of data covers a range between 165 ms and 550 ms, and we conclude them to be fixations of the nest region in the frontal visual field. We found no systematic relation between the locations of these fixations and the nest region: The flight sections, where the bumblebees keep the nest hole between -25° and 25° in their frontal visual field are distributed evenly across the entire area of the flights (fig. 8E).

**Fig. 8.**
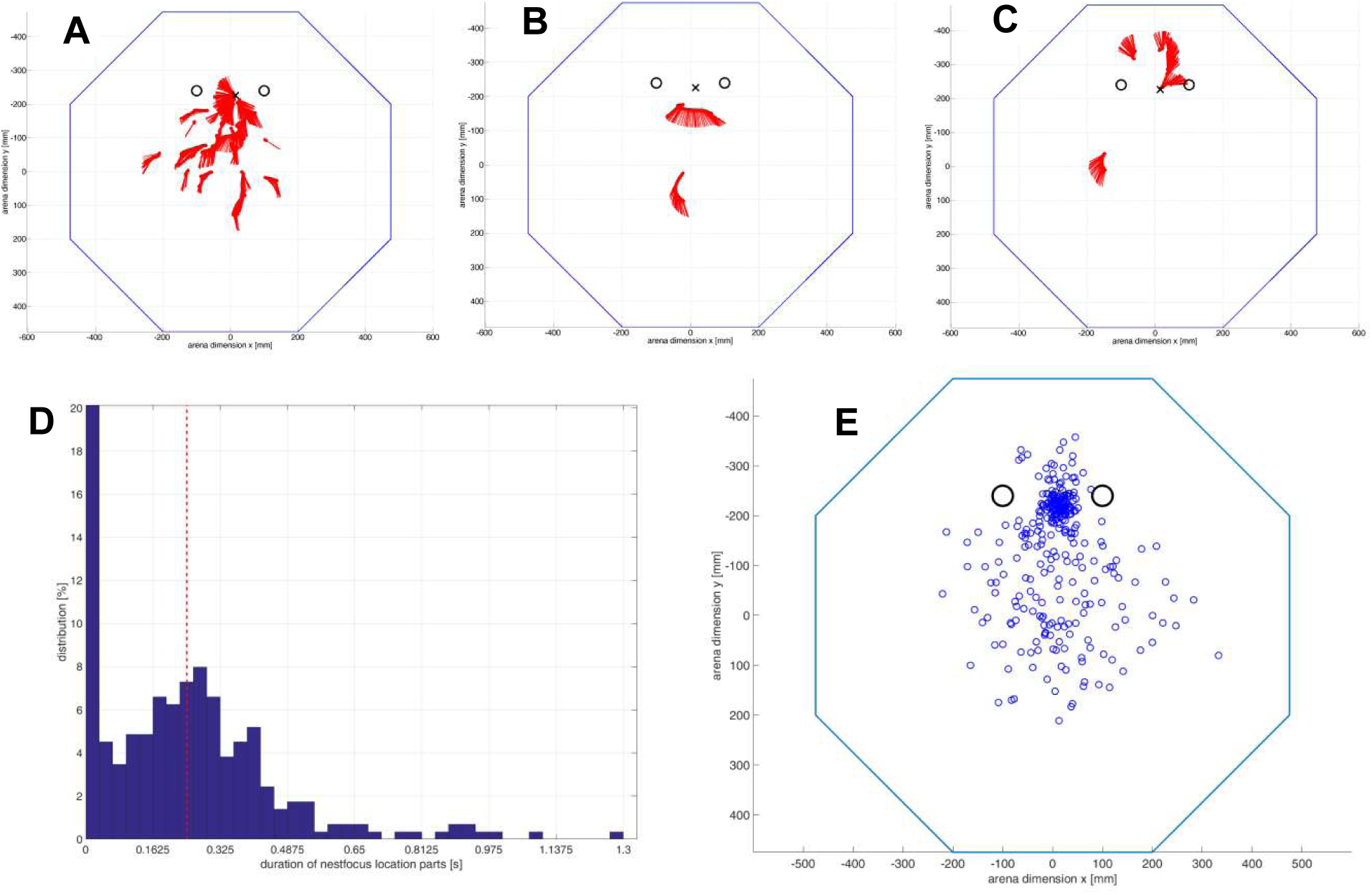
Locations in flight arena where bees fixate nest hole with frontal part of their visual field and duration of of fixations. (A-C) Locations of next fixations of the same first outbound flights and their duration as shown in Fig.2. Plotted is the position (red dots) and orientation (red lines) of the bumblebee in the arena when the nest hole is in the frontal visual field (between -25° and +25°). Time between consecutive dots: 20.27 ms. X: Nest hole, O: Cylinder, blue: arena walls. (D) Duration of individual nest fixations in seconds for all bumblebees in the 1^st^ flight phase (below 100 mm). n = 21 flights. (E) Locations in flight arena where bees fixate nest hole with frontal part of their visual field. Blue circles show the middle of each individual fixation section for all bumblebees in the 1^st^ flight phase (below 100 mm). Black circles: Cylinders, n = 21 flights.

### Sideward and forward components of flight

Flying insects, such as bees, perform a saccadic flight and gaze strategy to separate rapid head and body saccades from largely translational intersaccadic locomotion (Schilstra and van Hateren, 1999; Boedekker et al., 2010; Boeddeker et al., 2015; Braun et al., 2010; Braun et al., 2012; Geurten et al., 2010; Doussot et al., in prep). This strategy facilitates access to spatial information from the resulting optic flow, because only translational optic flow is distance dependent and contains spatial information (Egelhaaf et al., 2012).

A combination of pure translational and pure rotational movements in one flight segment, therefore, might be expected for outbound flights of bumblebees as well. Although, there are clear indications in our data for such a saccadic flight strategy (fig. 9A), the spatial resolution of our video footage was not sufficient given the chubby shape of bumblebees and the relatively large area that had to be filmed, to precisely address the temporal fine structure of the bees’ gaze strategy at the level of body orientation and, especially not at the level of head orientation. This issue will be tackled in detail in a forthcoming study (Doussot et al., in prep). Translational movements can be either forward/backward, sideward or a combination of both (diagonal) without changes in the yaw angle of the body orientation. To characterise the overall flight characteristic after leaving the nest hole and, especially, to what extent the bees performed sideward versus forward/backward movements, we determined the proportion of either of these components of translational movements. Flight sections, where sideward components are prevalent, are particularly relevant when spatial information is extracted from the retinal image flow in the frontal visual field, whereas forward or backward movement facilitates the extraction of spatial information in the lateral field. In the first flight phase sideward translational components predominated the flight pattern, while forward or backward movements were less prominent. This characteristic is specific for the initial phase of departure flights, as in later phases the proportion of sideward motion decreases over time and forward movements dominate the overall translatory flight component (fig. 9B-D). Flight manoeuvres with large sideways translational components close to a goal location are also known for honeybees (Dittmar et al., 2010; Braun et al., 2012) and hoverflies (Geurten et al., 2010). These sideways movements can be used by the insects to extract relative motion cues to estimate their distance to targets, such as the nest hole, which seems to be relevant in the early learning phase (Dittmar et al., 2010). These observations suggest that the sideward components during the initial phase of departure flights of bumblebees might play a role in gathering depth information in the close vicinity of the nest hole.

**Fig. 9.**
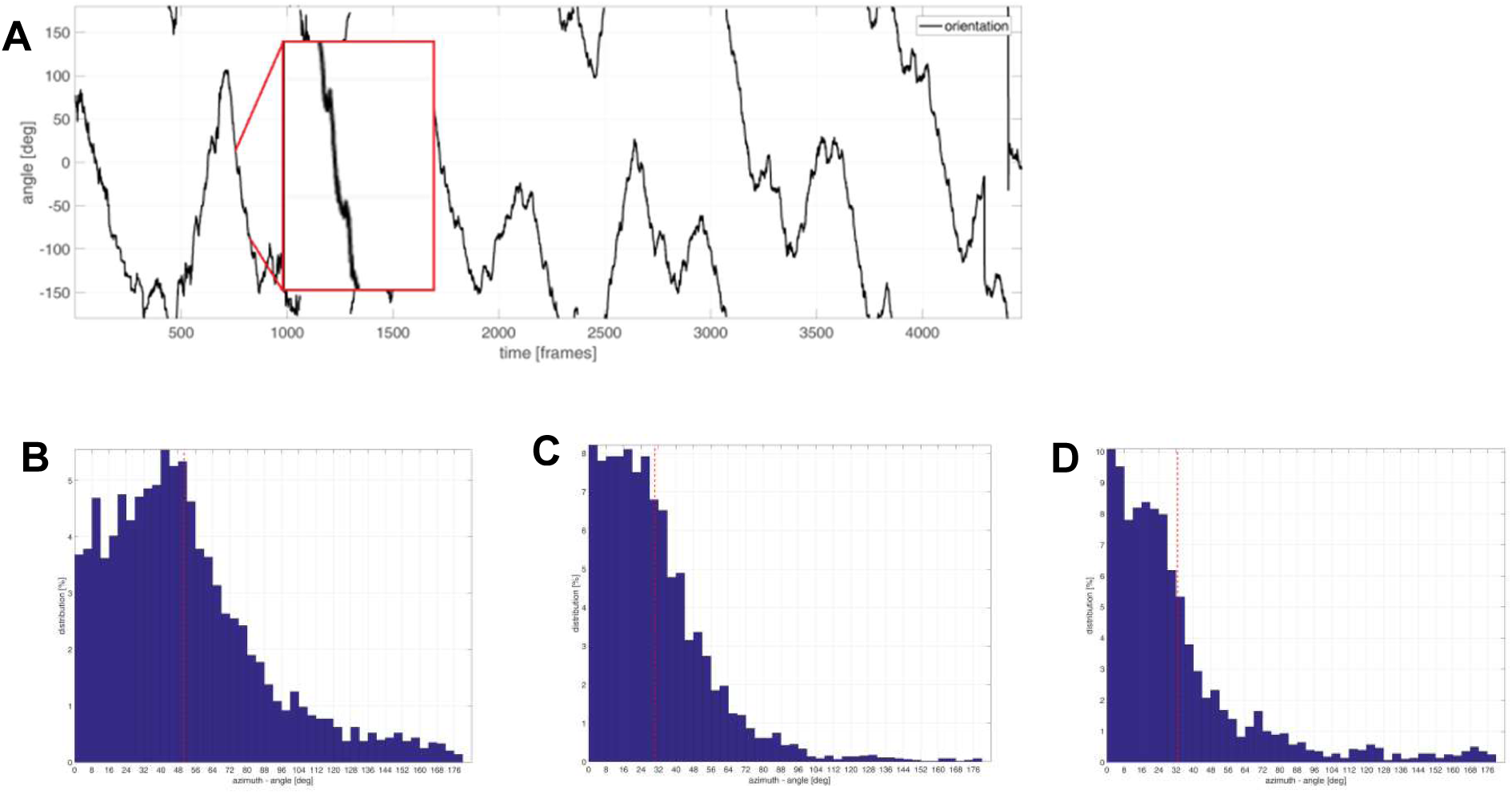
Saccadic flight structure. (A) Time course of body orientation of an example bumblebee on its 1^st^ departure flight. The red rectangle shows an inset of the orientation to show the characteristic saccadic flight structure in an enlarged fashion. (B) Sideward and forward components of flight: Distribution of direction of the translational component of motion relative to the orientation of the flight trajectory for all bumblebees for flight phase 1 (B), flight phase 2 (C) and flight phase 3 (D). The angle was determined from the ratio between the forward and sideward component of translation. The average angle is shown in red (dashed line: 50°, 32° and 33°). An angle of 0° is pure forward movement, an angle of 90° represents pure sideward movement.

### Changes in turn-direction (CTD) of the body

Not only translational movements play a role in an insect’s flight. During departure flights bumblebees perform loop-like excursions away from and back to the nest hole. Therefore, apart from translational flight sections, the flights show rotations of the bees’ body length axis (yaw rotations). Changes in turn-direction might be particularly relevant as they indicate decision points in flight behaviour. For social wasps such changes in turn-direction are generated at the end of the arcs characterising their departure flights and have been concluded to be elicited whenever the retinal image of the nest entrance moves to a lateral position in the visual field (Collett and Lehrer, 1993; Zeil et al., 1996; Zeil, 1993a; Zeil et al., 2007; Zeil et al., 2009). The changes of turn-direction, thus, lead to a correction of the accumulating retinal position error of the nest entrance (Zeil et al., 1993a).

Inspired by these observations we took a closer look at the changes in turn-direction of body orientation of bumblebees. The bees’ body orientation shows an alternating sequence of clockwise (cw) and counter-clockwise (ccw) rotations (fig. 10A). We analysed whether the reversals of turning direction are generated in specific spatial regions in the arena relative to the nest hole to get hints as to what environmental cues might trigger these changes. The locations where the bees perform CTD seem to be randomly distributed across the entire flight area during the initial phase of departure flights (fig. 10B). Nevertheless, we observed a tendency for more clockwise CTD when the nest hole was on the right side of the bee and more counter-clockwise CTD when the nest hole was on the left side (fig. 10Ci and 10Cii). This behaviour might reflect attempts of the bee to keep the nest hole region in the frontal visual field, performing a body rotation towards the nest when it leaves the fronto-lateral field. These attempts are performed in a similar, though not as precise way, as has been concluded for wasps (Zeil et al., 1996; Zeil, 1993a; Zeil et al., 2007; Zeil et al., 2009). This flight pattern disappears during later flight phases where the nest hole region might only play a minor role in shaping the flight (data not shown). For wasps Zeil (1993a) described a surprisingly constant rate of the CTD. For bumblebees we observed an average period of 1.6 s for the overall flight. Furthermore, we did not find any specific differences in the frequency of CTD for the different flight phases. Since the distance covered by the bee between CTD increased with altitude, the flight velocity during the turns increased accordingly (fig. 10D). This shows that bumblebees in our experiments seemed to have a specific frequency range, at which they performed the CTD. However, this range did not appear to be much affected by the bees’ position in the arena. Rather a CTD seemed to be initiated after a broadly constant time interval rather than a specific flight distance.

**Fig. 10.**
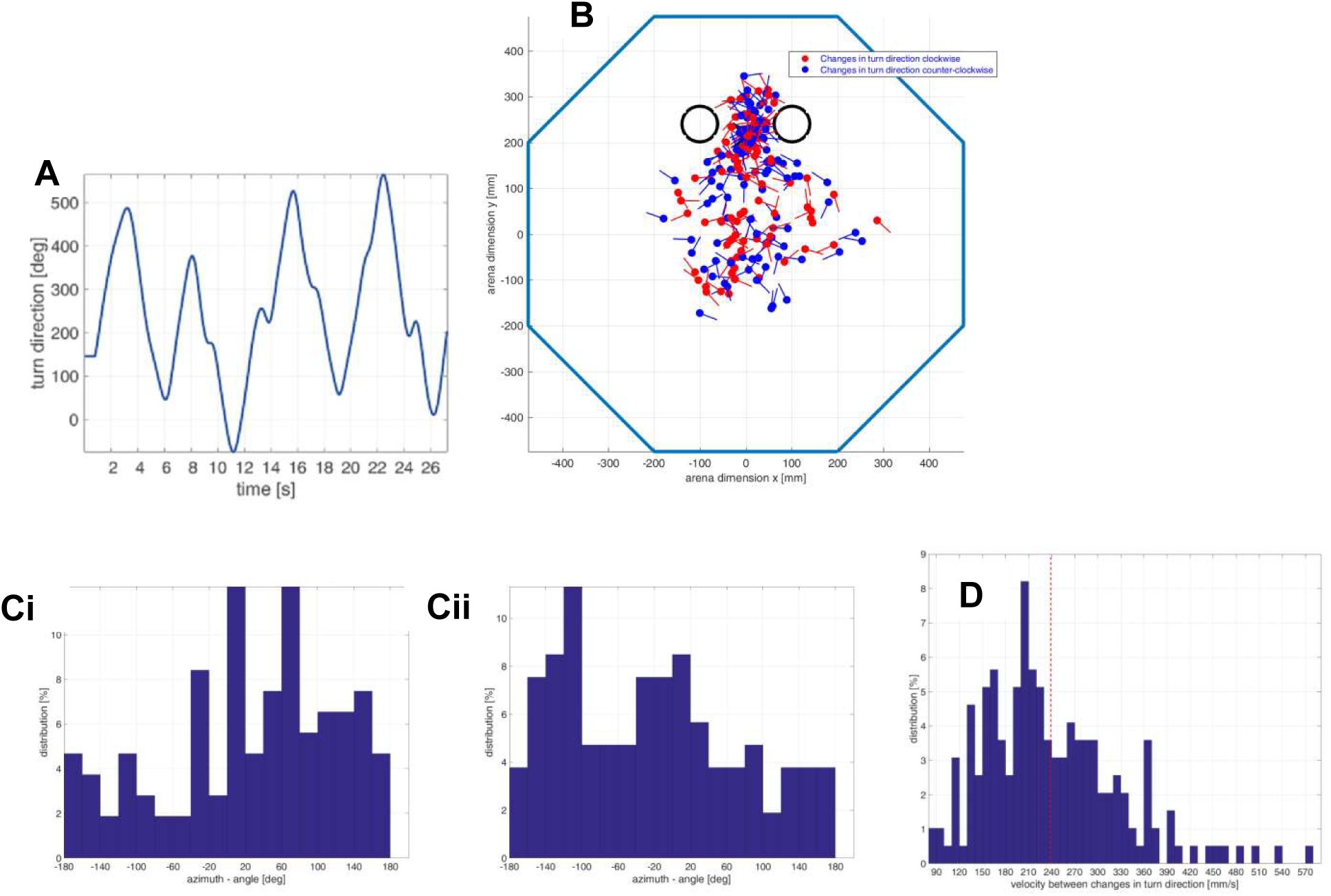
Changes in turn direction of body orientation. (A) Turn direction of body orientation of a bumblebee’s first departure flight as a function of time (B)Locations of changes in turn direction from clockwise to counter-clockwise and vice versa Clockwise and counter-clockwise turns for the 1^st^ departure flight of all bumblebees in phase 1 (below 100 mm). Black circles: Cylinders, n = 21 flights, bumblebee “architecture”: filled circle = head, line = body orientation (C) Retinal Position of the nest at clockwise and counter-clockwise changes in turn direction for the first outbound flight below 100 mm (i) clockwise, number of CDT = 106, (ii) counter-clockwise, number of CDT = 107, n = 21 flights (D) Flight velocity between changes in turn direction for the first outbound flight, below 100 mm, number of CDT = 195, dashed line: Mean of velocity, n = 21 flights

## Discussion

On their first outbound trip bumblebee foragers are confronted with unfamiliar and largely unpredictable surroundings of their nest hole. Therefore, they need to gather sufficient information about these surroundings before they leave the vicinity of the nest hole, in order to be able to find it again after a foraging trip. This implies a kind of innate learning program that controls at least the learning behaviour after a forager bee leaves the nest hole for the first time. The diversity of environments, however, makes it essential for the assumed innate learning program to be flexible to adjust it to the particular surroundings.

Previous studies propose that insects take some kind of panoramic information at the target location after leaving their nest. What information about the environment is stored and recalled on the return flights are still, to a large extent, open questions, as there is evidence for a wide range of possibilities. Representations about the environment might be based on a panoramic retinotopic snapshot of brightness values (Kollmeier et al., 2007) or of local motion values (‘ motion snapshot’; Dittmar et al., 2010). It might also be based on a more parsimonious representation, such as of the skyline of the horizon (Kollmeier et al., 2007; Philippides et al., 2011; Baddeley et al., 2011; Basten and Mallot, 2010; Wystrach et al., 2011; Graham and Cheng, 2009; Müller et al., in prep.). The information stored at the goal location is assumed to be compared in an appropriate way with the corresponding environmental information taken during the return flights to the nest. One way to accomplish this is to determine the similarity of retinotopic representations of the environment and to move in a way that increases the similarity (Cartwright and Collett, 1987; Vardy and Möller, 2005; Zeil et al., 2009). Another possibility is not to store the information on a retinotopic basis, but to determine an average landmark vector (ALV). The ALV is just the sum of vectors representing, for instance, the average brightness across elevation at each azimuthal position, or of the vectors pointing to ‘ landmarks’ identified in the retinal image.

Landmarks might be simple environmental features, such as trees. During the return flight the goal direction is determined according to this scheme at any location as a difference between the average landmark vector determined at the location and the corresponding vector determined at the current location (Lambrinos et al., 1999; Lambrinos et al., 2000). This kind of mechanism could be shown in model simulations to be sufficient to account, within a catchment area, for local homing, i.e. for returning of the agent back to its goal (Lambrinos et al., 2000; Möller, 2000; Stürzl and Mallot, 2006). The size and shape of the catchment area depends much on both the environment as well as the local homing mechanism.

All the mentioned models for the explanation of local homing in insects have in common that the information that is later used for returning to the goal is gathered locally at the goal location. These explanatory models, although they can explain local homing, seem to be somehow in disagreement with the concept of learning flights, where the insect is thought not to gather the relevant information just at the goal location, but during the often complex outbound flights after leaving the nest hole. That insects learn during their departure flights from the goal might be plausible, because of the animal’s heading direction during such flights: Wasps (Zeil, 1993a; Collett and Lehrer, 1993; Stürzl, 2016) honeybees (Lehrer, 1991; Lehrer, 1993; Lehrer and Collett, 1994; Dittmar, 2010; Dittmar, 2011) and bumblebees (Phillipides et al. 2013, Hempel de Ibarra et al. 2008; Riabinina et al., 2014; Collett et al., 2013) tend to orientate towards the goal location such as the nest hole or a food source for most or at least for the initial sections of their departure flights. This raises the question why insects should spend energy and time to perform a complex sequence of movements to gather information near their goal, if already one single panoramic snapshot taken at the goal location is sufficient for a successful return. This issue is further accentuated by the high degree of interindividual variability in the individual flight patterns of bumblebees as characterised here, but also between consecutive outbound flights of individual bees (Lobecke et al., in prep.), although there are obvious differences between different hymenopteran species in this regard (wasps: Zeil 1993a; Collett and Lehrer, 1993; honeybees: Lehrer and Collett, 1994).

The variability of outbound flights across bumblebees was systematically investigated in the present study: Although the overall flights differ tremendously between individuals, there are still common behavioural motifs in almost all outbound flights. Bumblebees leave the nest hole and spend the initial sections of departure close to the goal. They also roughly keep the nest hole region in their frontal visual field during periods in this initial section of the departure flights. Although the corresponding flight sections reveal a consistent spatial relationship to the nest hole and its vicinity, the are broadly spread in space in individual flights. After some time, the bees increase height and distance to their nest hole by performing loop-like manoeuvres. Thereby, the overall flight trajectories shift towards the centre of the flight arena. The nest hole region might play a role as a kind of trigger for changing the turn-direction, as has been proposed for solitary wasps (Zeil et al., 1993; Stürzl et al., 2009). However, the pattern of locations of changes in flight direction is highly variable in bumblebees: these locations may be almost everywhere in the area of the flight arena covered by the flight trajectories. Also the fine structure of the flights does not reveal obvious similarities between the bumblebees’ flight manoeuvres. Since the environment was kept constant in our experiments, this high variability can hardly be explained by the flexibility needed for such an innate behavioural learning program and the adaptivity of individuals to specific unpredictable environmental situations. Hence, the spatio-temporal structure of the departure flights of bumblebees does allow us to derive a consistent learning strategy.

The spatio-temporal characteristics of departure flights and, especially, the non-existence of a consistent pattern in their fine structure as well as the great interindividual variability led us to a new hypothesis regarding the functional significance of the departure flights. In particular, we hypothesise that bees gather information only during the very initial section of the flights, while they are still very close to the goal. In this section they are suggested to determine a dynamically representation of the surroundings as seen from a very small region around the goal (‘ semi-local dynamic snapshot’). The later flight sections of phase 1 of the departure flights (according to our classification explained in Results) are then hypothesised to be employed to probe the quality and usefulness of this semi-local information as well as the catchment area around the nest location. At the very beginning of the first outbound flight the initially naïve insect might gather information about the surroundings of the nest entrance only relatively locally, rather than during the entire phase 1 of the departure flights. Still, they might not take just a kind of panoramic stationary snapshot. Rather, bumblebees are assumed to have to move in the close vicinity of the nest hole: They need to turn around to get panoramic information about the environment. These rotations should be interspersed with at least brief translational flight intervals (e.g. intersaccadic intervals), if the animal needs to extract also information about the spatial layout of the environment from the perspective of the nest hole. All this information might then be combined to a semi-local representation of the behaviourally relevant environmental information quasi at the goal location. This information may then be employed as a basis of some local homing mechanism (see above). Further experiments are required which focus on the very initial phase of the departure flights while the bees move very close to the nest location; a high spatial resolution is then required to allow us resolving both body and head orientation in great detail. Since our current analysis covered the entire departure flights, this detailed analysis in not yet possible on this basis.

If our semi-local dynamic snapshot hypothesis were correct, the question is still open, why the bees do not just fly away after this very initial learning flight phase, but perform complex and time-consuming flight manoeuvers before eventually leaving the vicinity of the nest. We here further hypothesise that after this kind of semi-local dynamic snapshot has been determined at the goal location, the bee might test the reliability of this representation of the goal environment and especially the catchment area around the goal location. This might be a sensible strategy, because such a test phase might help to ensure that the learnt information about the nest location is really sufficient to be able to relocate it again after a foraging trip.

Overall, our hypotheses suggest that – in accordance with the common local homing models (see above) - semi-local information is sufficient to guide the insect back to its home location on the return flights. If this were correct, the observed inter-individual variability in the overall flight patterns would not be deleterious, because most of this part of the departure flights were not a component of a learning routine, but would just serve to probe the catchment area. This can, in principle, be done either systematically or by a somehow random procedure. This issue needs to be tested in further modelling analyses. In any case, as a consequence of such a scheme, the variability of departure flights is likely not to be the outcome of some kind of noise originating at any information processing stage in the nervous system, but part of a strategy probing the usefulness of the information acquired before at the goal location.

Upcoming studies investigating the initial learning behaviour in hymenopterans must be designed in a way to test, whether the phases after the initial sections of departure flights serve as a test for the reliability of the catchment area using semi-local dynamic information about the goal environment, actively gathered very close to the goal location.

## Acknowledgments

This study was supported by the Deutsche Forschungsgemeinschaft (DFG).

